# High prevalence of precocious menarche in Puerto Barrios, Guatemala

**DOI:** 10.1101/543090

**Authors:** Edmundo Torres-González, Griselda López, Britton Trabert, Hong Lou, Sarita Polo Guerra, Anali Orozco, Lisa Garland, Oscar Florez-Vargas, Miriam Castillo, Victor Argueta, Eduardo Gharzouzi, Michael Dean, Roberto Orozco

## Abstract

**Background:** Age of menarche is variable between women with a median age of 14 years old worldwide, and averages ranging from 12-13 years old in developed countries to 15-16 years old in low and middle-income countries. Precocious menarche, occurring before age 11, is rare, with a prevalence of 1.4 to 2.3% in most worldwide populations. Precocious menarche is poorly understood but is associated with early puberty and is a risk factor for pre-teen birth. In studying HPV prevalence in Latin America, we identified a community with a very high rate of precocious menarche.

**Objective(s):** Describe the patterns of precocious menarche in Guatemala.

**Study Design:** Reproductive histories were collected from 3385 cancer-free women at the time of routine Pap smear from 4 sites in Guatemala including hospitals in Guatemala City and Puerto Barrios, Izabal. Statistical analyses included determination of the age-specific prevalence of menarche and linear regression to determine the effect of year of birth, study site, number of births and miscarriages, on the age at menarche.

**Results:** Compared with a low prevalence of precocious menarche observed in Guatemala City (3.1%, 88/2834), we observed a high prevalence of precocious menarche in the city of Puerto Barrios, Izabal, Guatemala (88%, 486/551). We observed a high prevalence of precocious menarche in Puerto Barrios across all age groups. In contrast in Guatemala City, the median age at menarche declined from the age of 14 in 50-60-year-old women to 13 in women under 40 years of age. Hospital records show that the occurrence of both miscarriage and pregnancy under age 14 is substantially higher in Izabal. In addition, the main public hospital in Puerto Barrios accounts for a high fraction of the early pregnancies and miscarriages seen in Guatemala.

**Conclusions:** In Puerto Barrios, Izabal, Guatemala, the prevalence of menarche before age 11 is dramatically elevated compared to Guatemala City and substantially higher than other worldwide populations. We observed precocious menarche in Puerto Barrios in all age groups examined, indicating that this condition has been common for decades. This finding is supported by the comparatively higher occurrence of early pregnancy and miscarriage in Puerto Barrios compared with Guatemala City. The cause is unknown, but precocious menarche is associated with adverse reproductive outcome in young women and warrants further investigation.

## Introduction

The age of first menstruation or menarche is a defining reproductive milestone in women but is poorly understood. Biological, genetic and environmental factors influence the age at menarche^1^. Nutrition and physical/athletic activity are known to alter the age at menarche. Age at menarche varies considerably worldwide. In a meta-analysis of data from 67 countries, the mean age at menarche was 13.5 years (SD ± 0.98) with a range of 12.0-16.1 years^2^, and 13-14 years in Central America^2, 3^. Age at menarche has been observed to be higher in low and middle-income countries (LMICs) and is associated with calorie consumption (higher caloric consumption associated with earlier age at menarche)^2^.

Furthermore, many studies demonstrate that increased adiposity is associated with earlier onset of menarche^4–6^. An older age of menarche is observed in populations with a lower rate of literacy, and Thomas et al. suggest that this is due to the physical activity associated with child labor^2^. High levels of adolescent physical activity, particularly in female athletes, is associated with delayed menarche^7^.

Most countries with longitudinal data observe a decline in age at menarche over time^8–10^. For example, data from the US National Health and Nutrition Examination Survey (NHANES) demonstrates that the average age at menarche declined from 12.5 years in 1988–1994 to 12.3 years in 1999-2002^11^. Within the NHANES data, the age of menarche was highest in non-Hispanic whites (12.5) and lowest in Hispanics (12.2) and African Americans (12.0).

Precocious puberty is a complex condition and can be caused by abnormal production of sex hormones due to gonadal or adrenal tumors or exogenous exposures. Certain genetic conditions such as McCune–Albright syndrome or mutations in the *KIN28*, *LEP*, *LEPR*, and *MRKN3* genes are found in sporadic or familial cases. Also, childhood obesity is associated with earlier onset of female sexual development^12^. However, the condition is rare (1.4-2.3%) and is not related to a geographical location or socioeconomic status^11, 13–15^.

## Materials and Methods

Guatemala is the largest country in Central America with a population of approximately 16 million. The country is divided into 21 provinces known as Departamentos (Departments). The major population groups are indigenous Amerindians of the Mayan language groups; Europeans, predominantly from Spain and on the Caribbean coast; African populations such as the Garifuna^16^. Guatemala has a high cervical cancer incidence and mortality and a low rate of cervical cancer screening.

### Study populations

Women were recruited at three clinical centers (2 in Guatemala City, and Puerto Barrios in the Deptartment of Izabal) to participate in a study of the HPV prevalence in Guatemala. All healthy women (age 18-89) undergoing routine cytology screening at public clinics and hospitals were invited to participate. Women suspected of having cervical cancer were excluded. A total of 3385 (n=2834 in Guatemala City and 551 in the city of Puerto Barrios, Izabal) women were included as part of the HPV Prevalence in Guatemala Study^17^. Trained personnel administered a questionnaire on reproductive history and lifestyle factors^18^ to the women at the three sites: Insitituto de Cancerologia (INCAN), Guatemala City; Hospital General San Juan de Dios (HGSJDD), Guatemala City, and Hospital Nacional “Amistad Japón-Guatemala,” Puerto Barrios, Izabal, Guatemala. The questionnaire did not include questions on race or ethnic group.

The study was approved by the relevant Institutional Review Boards (IRB) in Guatemala and by the Office of Human Studies Research of the US National Institutes of Health and all subjects provided written informed consent.

We obtained information on births and miscarriages from specific departamentos and regional hospitals from the Guatemalan Ministry of Health http://epidemiologia.mspas.gob.gt and published sources^19, 20^.

### Statistical analyses

Statistical analyses were performed to determine the effect of year of birth, study site, the number of births and miscarriages, on the age at menarche using a binomial regression model (R version 3.5.0). A total of 5000 permutations were performed to estimate distribution of standardized regression coefficients for the overlap in observed beta coefficient calculated in the binomial logistic regression model. P < 0.05 was regarded as statistically significant.

## Results

In a study of HPV prevalence among women receiving cervical cancer screening, including 551 individuals in the city of Puerto Barrios, Guatemala and 2834 from Guatemala City (INCAN), we noticed a large discrepancy in the age at menarche. The participants’ mean age at enrollment was 37-45 years in Guatemala (**Table 1**). A total of 486 (88%, 486/551) of the women at Puerto Barrios reported menarche at age 10 or less, whereas only 3.1% (88/2834) of women at INCAN reported precocious menarche (**Figure 1**). The high proportion of precocious menarche in Puerto Barrios remained highly significant (p-value<0.0001) after controlling for age at enrollment, number of births, and miscarriages (**Table 2**). A permutation analysis demontrates that the precocious menarche in Puerto Barrios is unlikely to be due to chance; none of the resampled values produced a magnitude of the correlation as large as that observed (beta coefficient = 5.42).

**Table 1.** General characteristics of the samples

| **Hospital** | **INCAN** | **HGSJDD** | **HGSJDD** | **Hospital Nacional** |
| --- | --- | --- | --- | --- |
| **City** | Guatemala City | Guatemala City | Guatemala City | Puerto Barrios |
| **Location type** | Urban | Mixed | Urban | Urban |
| **Sample number** | 2618 | 145 | 71 | 551 |
| **Mean age (SD)** | 44.8 (10.1) | 40.8 (11.6) | 41.8 (11.0) | 36.6 (11.5) |
| **Median Age, *min-max*** | 44, 21-84 | 41, 19-87 | 41, 18-75 | 35, 17-79 |
| **Mean age at menarche (SD)** | 13.3 (1.6) | 13.1 (1.4) | 13.0 (1.6) | 9.8 (0.72) |
| **Median number of births, *min-max*** | 3, 0-20 | 3, 0-13 | 3, 0-11 | 2, 0-14 |
| **Median number of miscarriages, *min-max*** | 0, 0-6 | 0, 0-3 | 0, 0-3 | 0, 0-7 |

Data on the subjects from the four sample populations are shown. SD= standard deviation, Min=minimum, Max=maximum, INCAN, Instituto de Cancerologia; HGSJDD=Hospital General San Juan de Dios.

**Figure 1.**
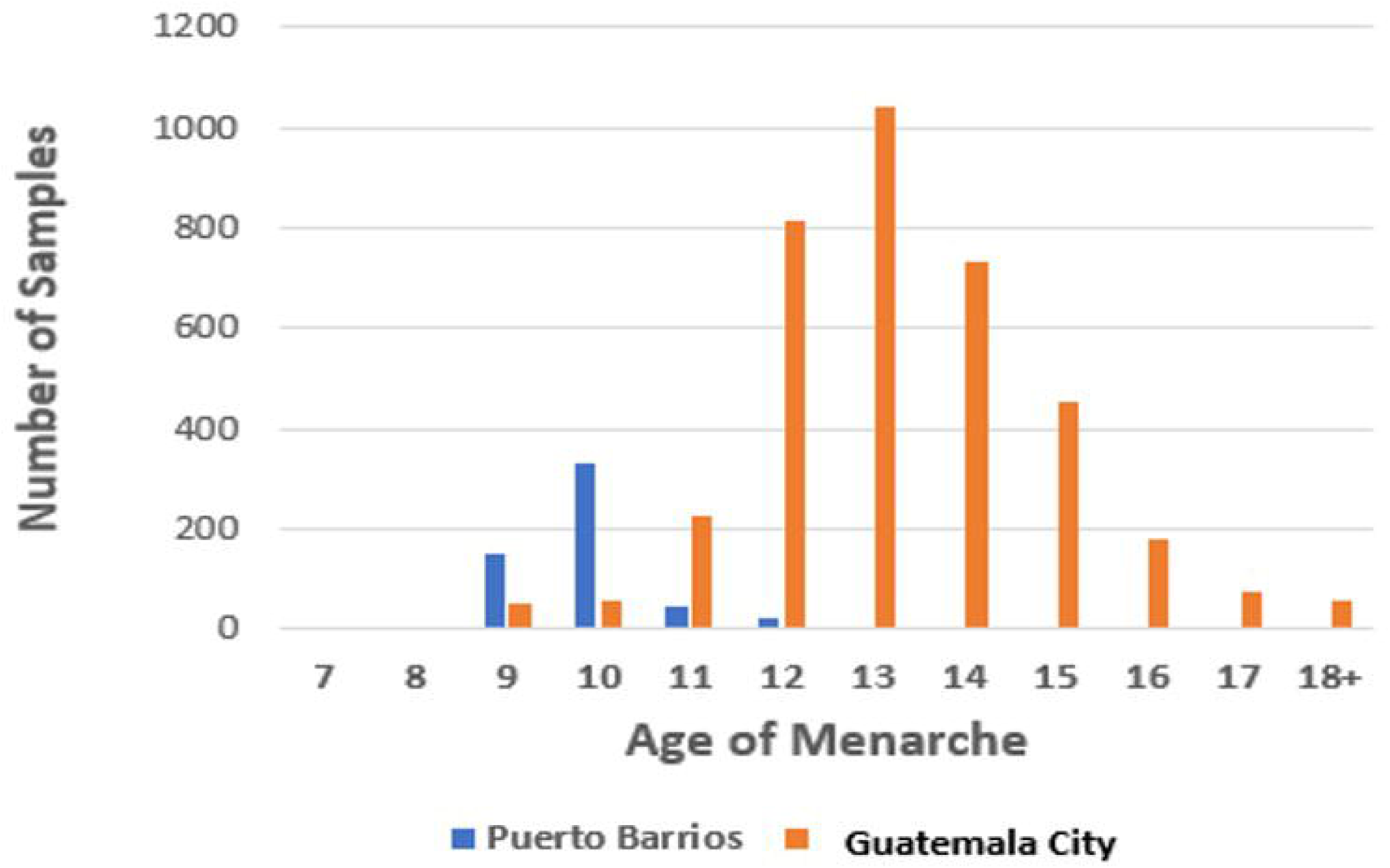
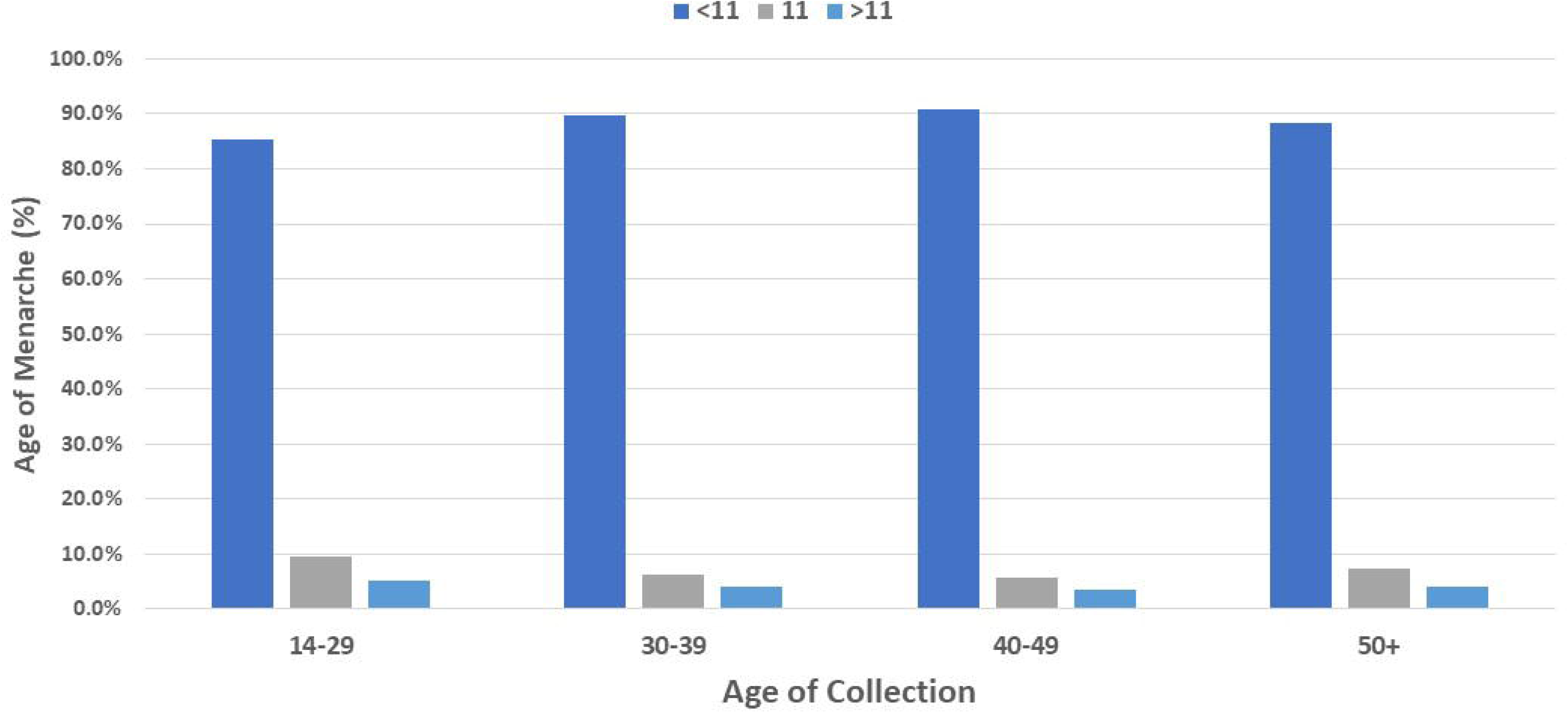
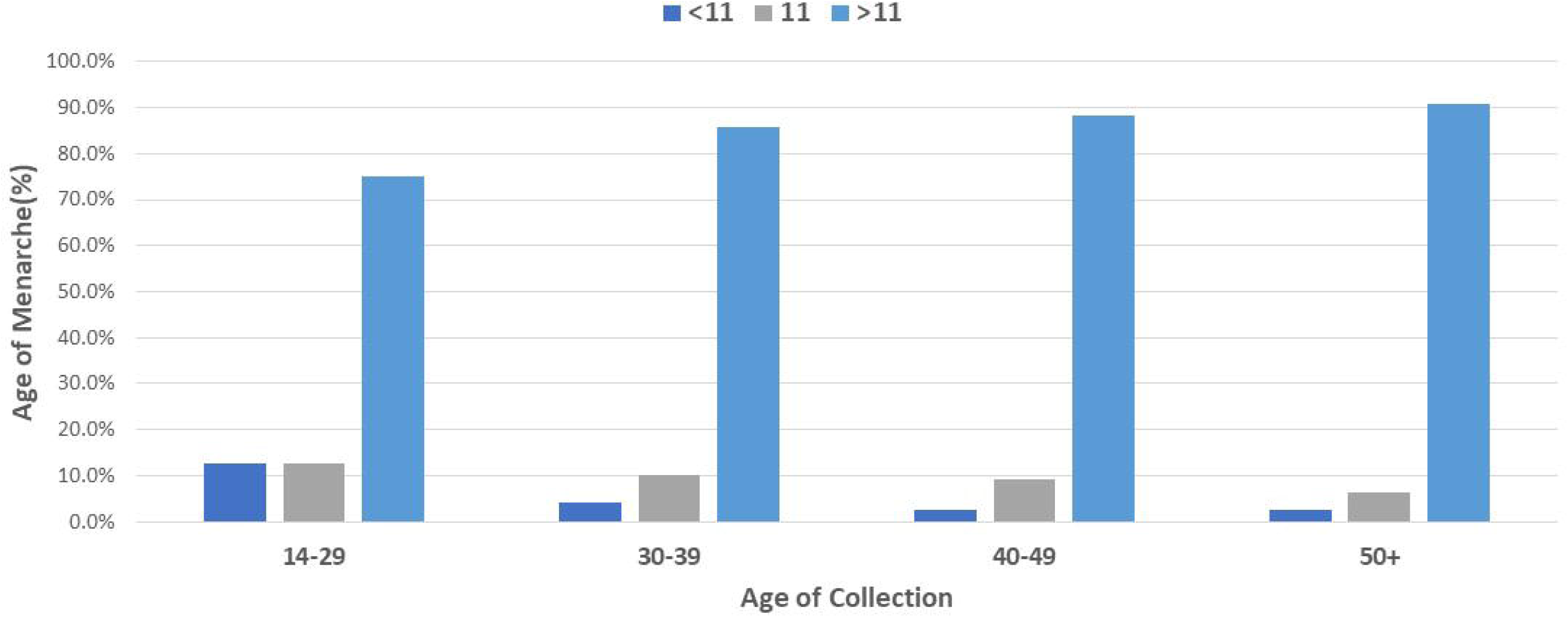
Age at menarche in women from Puerto Barrios (PB) and Guatemala City. A) the number of samples by age at menarche is shown for the two cities. B and C) The data is displayed by age group and by the age of menarche categories <age 11, age 11, and greater than 11 for Puerto Barrios (B) and Guatemala City (C).

**Table 2.** Regression coefficient and 95% confidence interval for the effect of year of birth, study site, number of births and miscarriages on precocious menarche (outcome).

| **Outcome:**<br>**Precocious Menarche** | **Model including age and study site** |  |  |
| --- | --- | --- | --- |
|  | **Regression**<br>**coefficient** | **95% CI** | **p value** |
| **Age at enrollment** | -0.011 | (-0.03, 0.01) | 0.26 |
| **Study site: Puerto Barrios** | 5.42 | (5.06, 5.81) | <0.001 *** |
| **No. births** | -0.024 | (-0.11, 0.06) | 0.60 |
| **No. miscarriages** | 0.010 | (-0.15, 0.37) | 0.44 |

Menarche was defined as a binary variable: Precocious menarche below age 11, Non-precocious menarche age 11 and above; and a binomial logistic model was used for regression. The data from both INCAN Guatemala City and Puerto Barrios were used concurrently in these models.

To determine if age at menarche is similar in different birth cohorts, we divided the women into four age groups as follows: 18-29, 30-39, 40-49, and ≥50 years old at enrollment. At Puerto Barrios, menarche before age 11 was consistently high (85-91%) in all age groups and was the highest in the 40-49 age group (91%). The median age at menarche was ten years old across all Puerto Barrios age groups. (**Figure 1B, C, Supplementary Table 1**). Therefore, there is an elevation of precocious menarche across all birth cohorts in Puerto Barrios. In Guatemala City, age at menarche <11 years old is 2-4% in the 30-39, 40-49 and ≥50 age groups. Nine out of 95 (9%) women in Guatemala City in the 18-29 year age group had menarche under age 11. (**Figure 1B, C).** The median age of menarche in Guatemala City was much higher than in Puerto Barrios, 13 in 18-29, 30-39, 40-49 age groups; increasing to 14 in the ≥50 age group (**Table 1**).

To evaluate adverse pregnancy outcomes associated with precocious menarche in Puerto Barrios and the Department of Izabal we accessed Ministry of Health data for all Deptartments of Guatemala. As displayed in **Figure 2**, Izabal accounts for 26% of all births in Guatemala in women under the age of 15, whereas this province has only 2.6% of the total births in the country. The hospital that is the recruitment site for the women of Puerto Barrios also has a high rate of births among females under the age of 15. This hospital reports 70% (7/10) of all births to girls 10 and 11 years of age reported among seven national hospitals, and 56% (31/55) of girls giving birth at age 12 (**Figure 3A**). There is a similarly high rate of miscarriage in girls age 10-14 in Puerto Barrios (**Figure 3B**).

**Figure 2.**
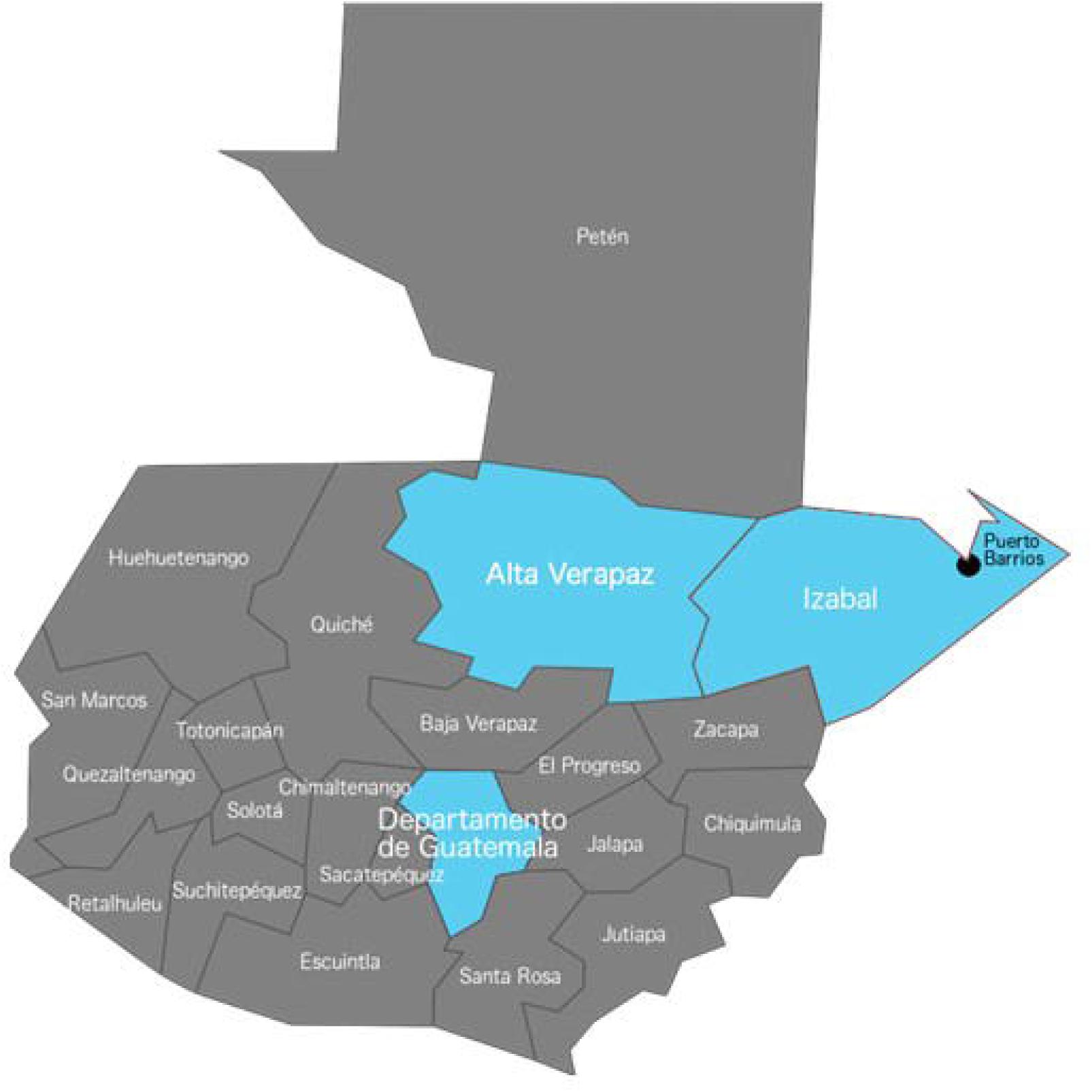
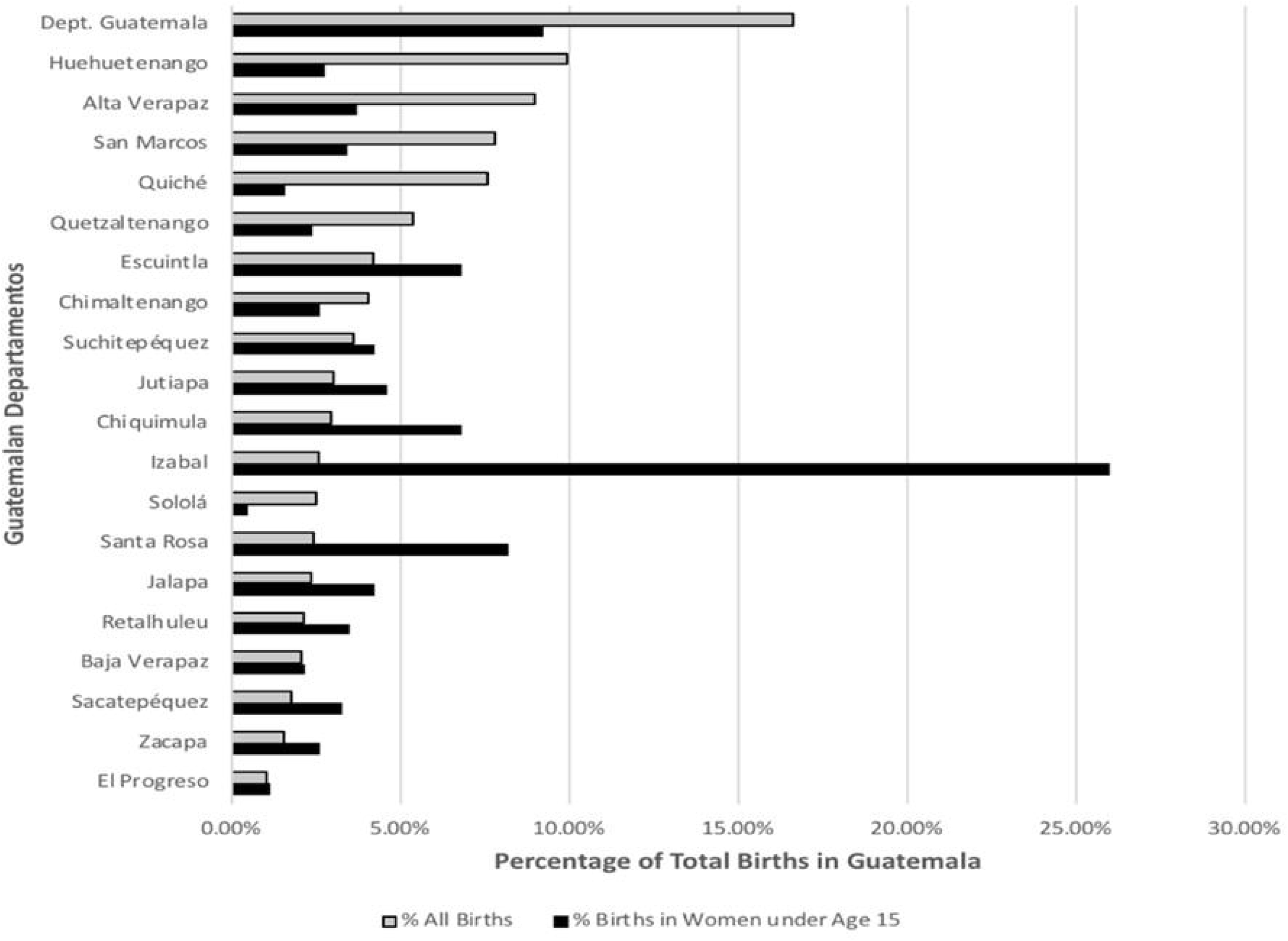
Map of the Departamentos of Guatemala and adolescent births. A), A map of Guatemala showing the location of the Departamentos of Izabal and Guatemala, and the collection sites. The location of Puerto Barrios, the site with elevated precocious menarche, is highlighted within Izabal. B) Births in Women under age 15 compared to all births in Guatemala (from Guatemalan Ministry of Health http://epidemiologia.mspas.gob.gt and published sources^19, 20^).

**Figure 3.**
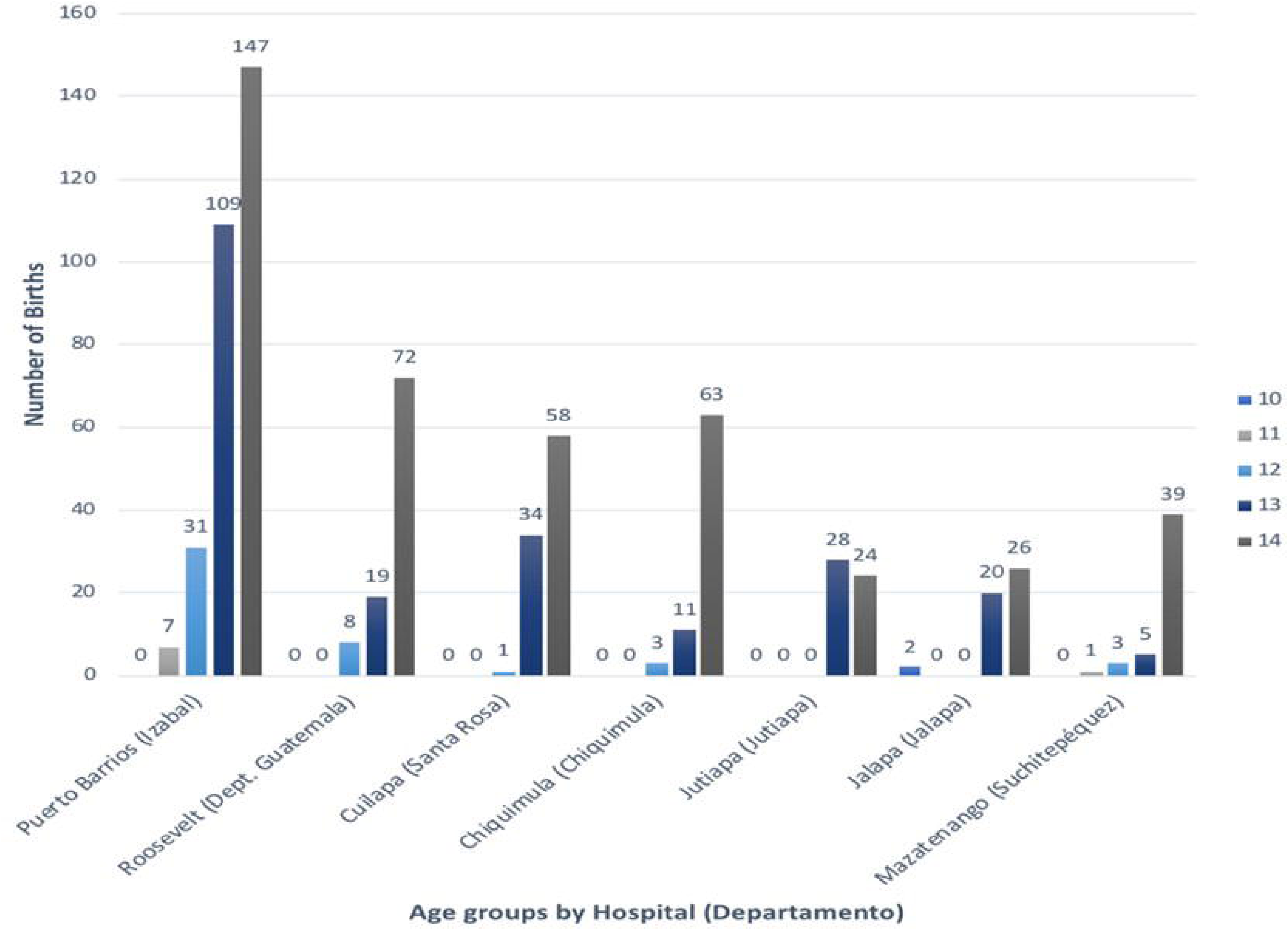
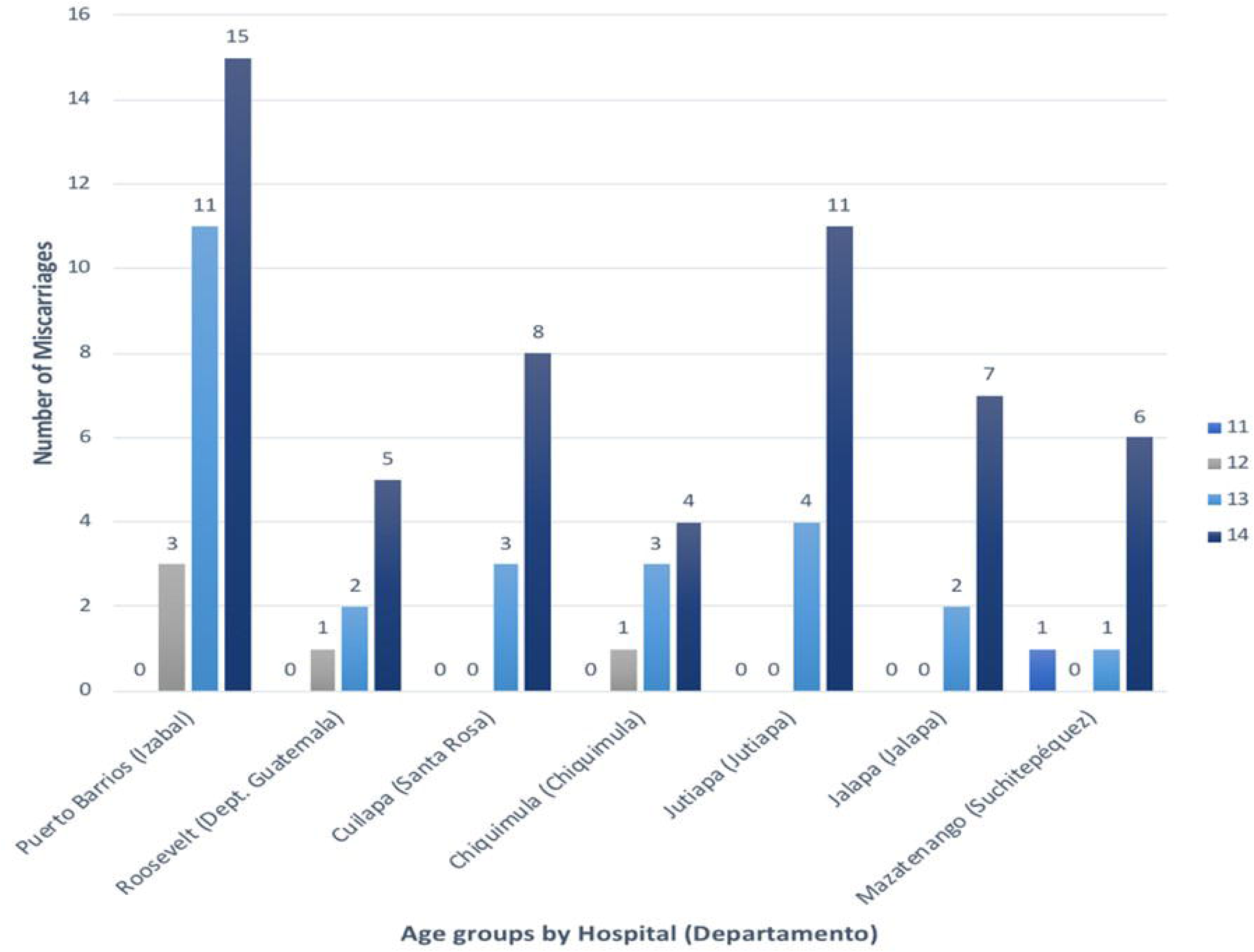
The number of births and miscariages in selected hospitals in Guatemala. A) Number of births to women under age 15 age for hospitals in Izabal, Guatemala City and four other Departaments. B) miscarriages for women age under age 15.

### Comment

In a survey of women receiving cytology screening at multiple hospitals in Guatemala, we identified a city (Puerto Barrios in the Department of Izabal) in Guatemala with an extraordinarilly high rate of precocious menarche, 88%. By contrast, precocious menarche is only 3.0% in subjects from Guatemala City, and we can find no reports of such a high prevalence of precocious menarche in general populations samples in any other region of the world. Comparison of birth year cohorts in Puerto Barrios and Guatemala City shows a consistently high rate of precocious menarche in 18-29, 30-39, 40-49 and over 50-year age groups, indicating that this phenomenon has been occurring in Puerto Barrios for a considerable period. Consistent with other world populations, age groups in Guatemala City show a steady decline in median age at menarche from 14 in women over age 50 to 13 in women under age 40^10^.

Evidence supporting early fertility in this region is demonstrated via national data in Guatemala in which 26% of the under 15 births occur in the Department of Izabal but only 2.6% of the total births. Furthermore, data from the same Puerto Barrios hospital show a high percentage of the births and miscarriage in 10-14-year-old females reported by national hospitals. This trend is continuing as data on registered births to adolescent women in Guatemala show that Izabal has an excess of births to women age 10-14 compared to Guatemala City or the rest of the country (data not shown, https://osarguatemala.org/embarazos-y-partos-de-madres-entre-10-y-19-anos-enero-a-junio-2018/).

Precocious puberty is a rare outcome in women worldwide^4, 21^ and is infrequently caused by genetic conditions, such as McCune Albright syndrome with mutations in the *GNAS* gene or mutations in the *KIN28*, *LEP*, *LEPR*, and *MRKN3* genes^12, 22^. Precocious puberty can also be due to abnormal production of sex hormones or to gonadal or adrenal tumors or hyperplasia^23, 24^. Also, childhood obesity is associated with earlier onset of female sexual development, but typically not a reduction in the age of menarche below 11 years^25^. Precocious menarche is a risk factor for early teen or pre-teen births and confers considerable morbidity and mortality. A study of 2 million women across Latin America shows that births in women age 15 and below have a four-fold higher risk of maternal death when compared to women age 20-24, and a 3.8-fold higher risk of puerperal endometritis (uterine infection)^26^.

We do not currently have detailed data on the reproductive or dietary history of our subjects or on circulating hormone levels. The population sample is from a public, urban hospital in a city of over 106,000 inhabitants, made up of admixed Amerindian/Europeans, Native Mayans (mostly speaking Q’eqchi’) and Garifuna, a population of African origin. Puerto Barrios is located on the Caribbean coast of Guatemala, adjacent to Honduras. A search of the literature does not reveal a population of either Caribbean, Central American, Garifuna or Maya with a similarly high prevalence of precocious menarche^2^. A genetic cause seems unlikely as this is not a homogenous population. Therefore, we suspect there is likely an environmental cause to the elevated level of precocious menarche. Puerto Barrios is geographically isolated from the rest of Guatemala, and it is possible there is a water contaminant in this region that may contribute to the earlier ages at menarche that we observed. The region has a warm, coastal climate and elevated rates of tropical, infectious, and sexually transmitted diseases. The area is also culturally unique with a strong influence of afro-Caribbean culture, and there may be a dietary explanation for this phenomenon. A potential environmental estrogenic toxin is zearalenone (ZEA), a mycotoxin produced by numerous species of *Fusarium* growing on corn and other food crops. Fusarium toxins have been found predominantly in the lowlands of Guatemala^27^, an area that would include the Department of Izabal, and are associated with precocious pubertal development^28^.

Our study has several limitations that would require a more in-depth analysis of age at menarche. We do not have detailed information on additional markers of female development such as age at thelarche, Tanner stage, or genetic ancestry. The sample is a convenience sample from a public hospital of women attending a cervical cancer screening clinic and may be biased by socioeconomic factors and language barriers. Our sample has an incomplete collection of menarche information. In total, 192 of 738 (26%) women in Puerto Barrios did not provide age at menarche data, whereas at INCAN, this rate is 413 of 3028 (1.7%) women. However, this missing data was not limited to a single age group and did not affect the overall trend in Puerto Barrios of elevated cases of precocious menarche; and a permutation analysis supports this conclusion. Strengths of the study include relatively large sample sizes and confirmatory data from independent databases.

In conclusion, we have identified a city in Guatemala that has a high rate of precocious menarche compared with the main city center (Guatemala City) as well as a high rate of births to women under 15 years of age. Understanding the factors contributing to precocious menarche in this population may be useful in helping to reduce adverse reproductive outcomes in young women in Guatemala.

## Acknowledgments

The authors would like to thank the staff and health professionals from the Instituto de Cancerología, Guatemala City, Guatemala, Hospital San Juan de Dios, Guatemala City, Guatemala and Patricia Zaid, Adolfo Santizo, Esther Avila and Lineth Boror for sample and data collection and shipping; the above individuals are employed by Biotec Guatemala, have no funding sources to disclose, and were not compensated separately for this study. We thank Christian Alvarez, Katherine McGlynn, and John Groopman for helpful discussions. Supported in part by the Intramural Research Program of the National Institutes of Health, National Cancer Institute, Center for Cancer Research, and from Leidos-Frederick under contract #HHSN261200800001E. The content of this publication does not necessarily reflect the views or policies of the Department of Health and Human Services, nor does mention of trade names, commercial products, or organizations imply endorsement by the U.S. government.

## Notes

The authors report no conflicts of interest.

## References

1. Marshall WA, Tanner JM. Variations in pattern of pubertal changes in girls. Arch Dis Child 1969;44:291–303.

2. Thomas F, Renaud F, Benefice E, De Meeus T, Guegan JF. International variability of ages at menarche and menopause: patterns and main determinants. Hum Biol 2001;73:271–90.

3. Khan AD, Schroeder DG, Martorell R, Haas JD, Rivera J. Early childhood determinants of age at menarche in rural guatemala. Am J Hum Biol 1996;8:717–23.

4. Kelly Y, Zilanawala A, Sacker A, Hiatt R, Viner R. Early puberty in 11-year-old girls: Millennium Cohort Study findings. Arch Dis Child 2017;102:232–37.

5. Harris MA, Prior JC, Koehoorn M. Age at menarche in the Canadian population: secular trends and relationship to adulthood BMI. J Adolesc Health 2008;43:548–54.

6. Leitao RB, Rodrigues LP, Neves L, Carvalho GS. Development of adiposity, obesity and age at menarche: an 8-year follow-up study in Portuguese schoolgirls. Int J Adolesc Med Health 2013;25:55–63.

7. Baker ER. Body weight and the initiation of puberty. Clin Obstet Gynecol 1985;28:573–9.

8. Graham MJ, Larsen U, Xu X. Secular trend in age at menarche in China: a case study of two rural counties in Anhui Province. J Biosoc Sci 1999;31:257–67.

9. Demerath EW, Towne B, Chumlea WC, et al. Recent decline in age at menarche: the Fels Longitudinal Study. Am J Hum Biol 2004;16:453–7.

10. Wyshak G, Frisch RE. Evidence for a secular trend in age of menarche. N Engl J Med 1982;306:1033–5.

11. Anderson SE, Must A. Interpreting the continued decline in the average age at menarche: results from two nationally representative surveys of U.S. girls studied 10 years apart. J Pediatr 2005;147:753–60.

12. Grandone A, Capristo C, Cirillo G, et al. Molecular Screening of MKRN3, DLK1, and KCNK9 Genes in Girls with Idiopathic Central Precocious Puberty. Horm Res Paediatr 2017;88:194–200.

13. Ameade EP, Garti HA. Age at Menarche and Factors that Influence It: A Study among Female University Students in Tamale, Northern Ghana. PLoS One 2016;11:e0155310.

14. Attallah NL. Age at menarche of schoolgirls in Egypt. Ann Hum Biol 1978;5:185–9.

15. Danker-Hopfe H, Delibalta K. Menarcheal age of Turkish girls in Bremen. Anthropol Anz 1990;48:1–14.

16. Martinez-Gonzalez LJ, Alvarez-Cubero MJ, Saiz M, Alvarez JC, Martinez-Labarga C, Lorente JA. Characterisation of genetic structure of the Mayan population in Guatemala by autosomal STR analysis. Ann Hum Biol 2016;43:457–68.

17. Lou H, Gharzouzi E, Guerra SP, et al. Low-cost HPV testing and the prevalence of cervical infection in asymptomatic populations in Guatemala. BMC Cancer 2018;18:562.

18. Lou H, Villagran G, Boland JF, et al. Genome Analysis of Latin American Cervical Cancer: Frequent Activation of the PIK3CA Pathway. Clin Cancer Res 2015;21:5360–70.

19. Alvarez T. Atencion Hospitalaria del Parto y Aborto en Adolescentes Embarazadas antes de los 14 Anos. Guatemala City: Universidad San Carlos de Guatemala, 2013.

20. Orozco Vasquez I. Edad de la menarquia en la poblacion Guatemalteco. Guatemala City: Universidad de San Carlos de Guatemala 1999.

21. Le MOAL J, Rigou A, Le Tertre A, De Crouy-Channel P, Leger J, Carel JC. Marked geographic patterns in the incidence of idiopathic central precocious puberty: a nationwide study in France. Eur J Endocrinol 2018;178:33–41.

22. Robinson C, Collins MT, Boyce AM. Fibrous Dysplasia/McCune-Albright Syndrome: Clinical and Translational Perspectives. Curr Osteoporos Rep 2016;14:178–86.

23. Carel JC, Leger J. Clinical practice. Precocious puberty. The New England journal of medicine 2008;358:2366–77.

24. Parent AS, Teilmann G, Juul A, Skakkebaek NE, Toppari J, Bourguignon JP. The timing of normal puberty and the age limits of sexual precocity: variations around the world, secular trends, and changes after migration. Endocr Rev 2003;24:668–93.

25. Biro FM, Wien M. Childhood obesity and adult morbidities. Am J Clin Nutr 2010;91:1499S–505S.

26. Cavazos-Rehg PA, Krauss MJ, Spitznagel EL, et al. Maternal age and risk of labor and delivery complications. Matern Child Health J 2015;19:1202–11.

27. Torres OA, Palencia E, Lopez De Pratdesaba L, et al. Estimated fumonisin exposure in Guatemala is greatest in consumers of lowland maize. J Nutr 2007;137:2723–9.

28. Massart F, Saggese G. Oestrogenic mycotoxin exposures and precocious pubertal development. Int J Androl 2010;33:369–76.

