## Supplemental Table 1 for "High prevalence of precocious menarche in Puerto Barrios, Guatemala"

**Supplemental Table 1.** Raw numbers for age at menarche by age groups in INCAN and PB

| Puerto Barrios | Age at Menarche (AaM) | | | | | | | | | | | | | | | | | | |
|---|---|---|---|---|---|---|---|---|---|---|---|---|---|---|---|---|---|---|---|
| **Age at Collection (AaC)** | **8-9** | | **10** | | **11** | | **12** | | **13** | | **14** | | **15** | | **16+** | | **Total** | **Median AoM** | **Mean AoM** |
| **18-29** | 46 | 26% | 106 | 60% | 17 | 10% | 9 | 5% | 0 | 0% | 0 | 0% | 0 | 0% | 0 | 0% | 178 | 10 | 9.93 |
| **30-39** | 39 | 22% | 118 | 67% | 11 | 6% | 5 | 3% | 1 | 1% | 1 | 1% | 0 | 0% | 0 | 0% | 175 | 10 | 9.9 |
| **40-49** | 41 | 29% | 89 | 62% | 8 | 6% | 5 | 3% | 0 | 0% | 0 | 0% | 0 | 0% | 0 | 0% | 143 | 10 | 9.83 |
| **50+** | 30 | 32% | 54 | 57% | 7 | 7% | 2 | 2% | 1 | 1% | 0 | 0% | 1 | 1% | 0 | 0% | 95 | 10 | 9.89 |
| **Total No.** | 156 | | 367 | | 43 | | 21 | | 2 | | 1 | | 1 | | 0 | | 591 | | |

| Guatemala City | Age at Menarche (AaM) | | | | | | | | | | | | | | | | | | |
|---|---|---|---|---|---|---|---|---|---|---|---|---|---|---|---|---|---|---|---|
| **Age at Collection (AaC)** | **8-9** | | **10** | | **11** | | **12** | | **13** | | **14** | | **15** | | **16+** | | **Total** | **Median AaM** | **Mean AaM** |
| **18-29** | 4 | 17% | 5 | 21% | 6 | 25% | 30 | 125% | 24 | 100% | 12 | 50% | 11 | 46% | 3 | 13% | 24 | 13 | 12.9 |
| **30-39** | 8 | 1% | 28 | 4% | 85 | 12% | 195 | 27% | 252 | 34% | 177 | 24% | 103 | 14% | 60 | 8% | 731 | 13 | 13.1 |
| **40-49** | 9 | 1% | 13 | 2% | 75 | 11% | 176 | 25% | 147 | 21% | 183 | 26% | 140 | 20% | 79 | 11% | 699 | 13 | 13.36 |
| **50+** | 6 | 1% | 13 | 3% | 46 | 9% | 164 | 32% | 197 | 38% | 211 | 41% | 126 | 24% | 100 | 19% | 519 | 14 | 13.61 |
| **Total No.** | 27 | | 59 | | 212 | | 565 | | 620 | | 583 | | 380 | | 242 | | 1973 | | |
